# Community structure – ecosystem function relationships in the Congo Basin methane cycle depend on the physiological scale of function

**DOI:** 10.1101/639989

**Authors:** Kyle M. Meyer, Anya M. Hopple, Ann M. Klein, Andrew H. Morris, Scott Bridgham, Brendan J. M. Bohannan

**Author notes:** Authors contributed equally to the preparation of this manuscript (Kyle M. Meyer, Anya M. Hopple). Department of Integrative Biology, University of California – Berkeley, Berkeley, CA, USA.

## Abstract

Belowground ecosystem processes can be highly variable and difficult to predict using microbial community data. Here we argue that this stems from at least three issues: 1) complex covariance structure of samples (with environmental conditions or spatial proximity) can make distinguishing biotic drivers a challenge, 2) communities can control ecosystem processes through multiple mechanisms, making the identification of these controls a challenge and 3) ecosystem function assessments can be broad in physiological scale, encapsulating multiple processes with unique microbially mediated controls. We test these assertions using methane (CH_4_)-cycling processes in soil samples collected along a wetland-to-upland habitat gradient in the Congo Basin. We perform our measurements of function under controlled laboratory conditions and include environmental covariates in statistical analyses to aid in identifying biotic drivers. We divide measurements of microbial communities into four attributes (abundance, activity, composition, and diversity) that represent different forms of community control. Lastly, our process measurements differ in physiological scale, including broader processes (gross methanogenesis and methanotrophy) that involve more mediating groups, to finer processes (hydrogenotrophic methanogenesis and high-affinity CH_4_ oxidation) with fewer mediating groups. We observed that finer scale processes can be more readily predicted from microbial community structure than broader scale processes. In addition, the nature of those relationships differed, with broad processes limited by abundance while fine-scale processes were associated with diversity and composition. These findings demonstrate the importance of carefully defining the physiological scale of ecosystem function and performing community measurements that represent the range of possible controls on ecosystem processes.

## INTRODUCTION

Belowground ecosystem processes can be highly variable, and there have been numerous attempts to explain or predict this variation using microbial community data. However, our ability to identify microbial community drivers of processes remains in its infancy. This challenge stems from at least three issues. First, community variables often have complex covariance structure (e.g. spatial or environmental autocorrelation) (Legendre, 1993), which can complicate the task of distinguishing the biotic signal underlying ecosystem process rate variation. Second, communities can control processes through a number of mechanisms, making the identification of these controls a challenge (Bier et al., 2015). For example, communities can influence the rate of a process through abundance (of particular groups or the community as a whole), composition (i.e. the identity of community members), diversity, and activity, each of which would require the quantification of different community attributes. Lastly, function is often assessed at a broad physiological scale that encapsulates many processes that could have unique controls (Inkpen et al., 2017). For example, gross decomposition of organic matter or net flux of methane (CH_4_) into the atmosphere are commonly measured belowground ecosystem functions, yet these measurements represent the aggregate result of many different physiological processes.

We tested this idea using the two counteracting microbial processes that largely control atmospheric CH_4_ emissions from soils: CH_4_ production (methanogenesis) and CH_4_ consumption (methanotrophy). Each of these processes can be further divided into underlying processes that involve fewer mediating microbial taxa. For instance, gross methanogenesis can have direct inputs from at least three processes: hydrogenotrophic (which uses CO_2_ and H_2_ to produce CH_4_), acetoclastic (which uses acetate), and/or methylotrophic (which uses methanol, methyl-amides, or methyl-sulfides) methanogenesis. The process of methanotrophy can be subdivided as well. Methanotrophy at high CH_4_ concentrations (typical of those found in waterlogged soil and sediment) is performed by a wide diversity of methanotrophs found in the Gammaproteobacteria, Alphaproteobacteria, and Verrucomicrobia (Knief, 2015), whereas the consumption of CH_4_ at atmospheric concentrations (a.k.a. high-affinity CH_4_ oxidation) appears to be much more phylogenetically restricted and is likely performed by many fewer taxa (Ho et al., 2013; Knief, 2015; Kolb, 2009). Taken together, these observations suggest that gross measurements of either methanogenesis or methanotrophy may mask subtle variations in these underlying processes and could thereby blur the connection between community variation and ecosystem function.

A major challenge in the study of ecosystem function lies in identifying underlying community controls. Group abundance is one such form of control, whereby the rate of a process only proceeds as quickly as the total number of capable individuals. There are examples of this control for both methanogenesis (Ma, Conrad, & Lu, 2012) and methanotrophy (Freitag & Prosser, 2009), but many studies have found no such relationship (see Rocca et al., 2014). While this could be driven in part by technical issues (e.g. abundance being inaccurately assessed due to primer bias), processes may be controlled by other community attributes. Most soil organisms are in an inactive state at any given time (Lennon and Jones, 2011). Low activity levels can limit the rate of ecosystem processes either by decreasing syntrophic interactions or preventing substrate uptake. Measurements of methanotroph and methanogen transcriptional activity levels have been shown to positively correlate with process rates (Freitag & Prosser, 2009; Freitag, Toet, Ineson, & Prosser, 2010), suggesting that the activity levels of key groups can constrain these processes. Moreover, community assessments using RNA may more closely approximate the active fraction of the community (Jones & Lennon, 2010; Kamke et al., 2010; Baldrian et al., 2012; although see Papp et al., 2018a) and therefore provide a means to approximate activity-based community controls.

The composition of a community can impose another constraint on ecosystem processes. Composition control (a.k.a. species identity effects (Bannar-Martin et al., 2018)) could occur if certain community members perform a process more efficiently or rapidly than others, and those taxa/traits are dispersal limited. In this case, certain individuals would be expected to more strongly correlate with process rate variation than others. Distinguishing such a compositional signal can be a challenge because communities tend to co-vary depending on their spatial proximity or similarity in environmental controls (Legendre, 1993). There is precedent to suggest that community composition is an important determinant of CH_4_-cycling rates. For instance, even closely related methanogen/methanotroph taxa can differ in traits such as substrate affinity, substrate preference, environmental tolerance, or competitive ability (Garcia, Patel, & Ollivier, 2000; Ho et al., 2013; Knief, 2015; Nazaries, Murrell, Millard, Baggs, & Singh, 2013). Therefore, differences in community composition have the potential to influence CH_4_-cycling processes, which has been shown in several systems (Bodelier et al., 2013; McCalley et al., 2014; Nazaries, Pan, et al., 2013; Sierocinski et al., 2017, 2018). Membership of other functional guilds within a community is also an important consideration, particularly for processes that require metabolic byproducts as the primary substrate. Methanogens depend on the fermentative byproducts (*e.g.* H^+^ and acetate) of other community members and must compete for these substrates with other community members. This is also the case for methanotrophs, where competition for O_2_ and/or other soil nutrients can limit rates of methanotrophy (Bodelier, Roslev, Henckel, & Frenzel, 2000). Thus, *who is there* may be a particularly important consideration for predicting the rates of ecosystem processes, particularly if efficient traits are dispersal limited or competition for substrates is common.

Lastly, a growing body of work suggests that biodiversity may affect rates of ecosystem processes (Tilman, Wedin, & Knops, 1996), as well as the resistance, resilience, or stability of a process through time (Loreau et al., 2001; Tilman & Downing, 1994). This pattern could arise through multiple ecological mechanisms including: 1) niche complementarity, 2) sampling effects, or 3) facilitation. It has been argued that ecosystem functions mediated by a wide assortment of taxa may be less influenced by levels of diversity than those mediated by relatively fewer taxa (Schimel, 1995; Schimel & Gulledge, 1998). This would suggest that as one narrows the physiological scale of ecosystem function, the diversity of participants should become a stronger determinant of process rates. The abilities to produce and consume CH_4_ are both highly phylogenetically conserved (Martiny, Treseder, & Pusch, 2013), and relatively few taxa possess the ability to carry out these processes. There are even fewer taxa that perform underlying processes such as hydrogenotrophic methanogenesis and high-affinity methanotrophy (Knief, 2015; Kolb, 2009). Positive diversity – ecosystem function relationships have been observed for methanotrophy experimentally using intact soil cores from the field (Bodelier et al., 2013), as well as in artificially assembled methanotroph communities (Schnyder, Bodelier, Hartmann, Henneberger, & Niklaus, 2018). This has also been observed for methanogenesis in anaerobic digesters (Sierocinski et al., 2018). Although several studies have tested whether diversity and process rates are related, few have asked whether adjusting the physiological scale at which function is assessed improves this relationship.

Here we present results from a set of experiments designed to identify the biological drivers of variation in CH_4_-cycling dynamics. Our goals were to determine (1) whether we could identify significant community controls on CH_4_ cycling after accounting for covariance structure, and (2) whether the strength and nature of those controls changes with the physiological scale of the process. To do this, we collected soils along a diverse wetland-to-upland habitat gradient in the Congo Basin, a region that has received very little scientific attention for its role in the global CH_4_ cycle (Bridgham, Cadillo-Quiroz, Keller, & Zhuang, 2013; Kirschke et al., 2013) and has been shown to exhibit highly dynamic spatial and temporal CH_4_ cycling dynamics (R. A. Delmas, Tathy, & Cros, 1992; R. Delmas, Servant, Tathy, Cros, & Labat, 1992; MacDonald et al., 1999; Tathy et al., 1992). We estimate process rate potentials under controlled laboratory conditions, which minimizes the impact of environmental variation and enhances the biotic signal underlying process rate variation. We varied the physiological scale at which we assessed ecosystem function, ranging from broader scales including gross methanogenesis and methanotrophy, to finer scales including hydrogenotrophic methanogenesis and high-affinity methanotrophy. We paired RNA- and DNA-based microbial community measurements with each process measurement and calculated four sets of community attributes (abundance, activity, composition, and diversity) to aid in the identification of putative community controls. We hypothesized that there would be a connection between microbial community structure and CH_4_-cycling processes after accounting for environmental variation, and that finer scale processes *(i.e.* those involving a lower diversity of mediating taxa) would exhibit a stronger relationship with community attributes than broader processes, as exemplified by a higher proportion of variance explained. Our results suggest that communities are an important driver of CH_4_-cycling processes in tropical ecosystems, and that the strength and nature of the relationship between community structure and ecosystem function can depend on the physiological scale by which function is assessed.

## METHODS AND MATERIALS

### Site selection and sampling

We selected a diverse set of 15 sites in Southwestern Gabon in and around the Gamba Complex of Protected Areas (Lee, Alonso, Dallmeier, Campbell, & Pauwels, 2006). Gabonese ecosystems belong to the Guineo-Congolian regional center of endemism (White, 1979), making them broadly representative of the Congo Basin. Our sites included 4 mineral-soil forested wetlands, 2 peatlands, 2 seasonally wet mineral-soil forests, 3 upland grasslands (2 with termite mounds), 2 upland forests, a 1-year old plantation, and 1 abandoned plantation. Seasonal wetlands were identified by tree species composition, thick O and/or A soil horizons with mottling, and tree watermarks. Water tables were at least 40 cm below the soil surface in seasonal wetlands. Our sampling took place at the beginning of the Gabonese rainy season, from October to November 2014.

### Potential gross and high-affinity CH_4_ oxidation rates

We collected three intact soil cores from above the water table at the beginning, middle, and end of 35 m field transects established at each site. The samples were stored in PVC tubes (5 cm dia., 8 cm height) to maintain soil structure and kept at *in situ* air temperatures (28 °C) until the end of the sampling trip (~3 weeks). One week after returning to the University of Oregon, each PVC core was placed into a gas-tight Mason jar that was retro-fitted with a headspace sampling port and incubated at 28 °C in the dark. Rates of aerobic CH_4_ oxidation were determined under initially low (5 ppm CH_4_/mL in headspace) and high (1000 ppm CH_4_/mL headspace) CH_4_ concentrations, to estimate rates of high-affinity and gross methanotrophy, respectively. CH_4_ concentration measurements were taken at 0.33, 3, 6, and 9 hours for high-affinity methanotrophy and at 0.33, 3, 6, 9, 24, and 48 hours for gross methanotrophy. The same soil cores were used to determine high-affinity methane oxidation and gross methanotrophy in quick succession. We applied a first-order exponential decay function to determine the rate constant (k, units = d^-1^; *i.e.* dCH_4_/dt = k[CH_4_]) of the exponential decrease in CH_4_ when at low initial CH_4_ concentrations (i.e. high-affinity methanotrophy). At high CH_4_ concentrations we measured the linear decreases in CH_4_ to determine the maximum velocity rates (V_max_, units = μg CH_4_ cm^-3^ d^-1^) (i.e. gross methanotrophy). During our high-affinity CH_4_ oxidation incubations, CH_4_ concentrations exponentially decreased in all samples, except for one, across our four measurements; thus, all time points were included in subsequent calculations. While one sample did not immediately exhibit CH_4_ oxidation at the first timepoint (0.33 hours), it did show exponential decreases in CH_4_ concentrations following the 3 hour timepoint; thus, only three timepoints were used to calculate the k-value for this sample. During our gross methanotrophy incubations, we included only timepoints that encompassed the linear portion of the exponential decrease in CH_4_ concentration, which ranged from 3-5 timepoints depending on the sample.

### Potential gross methanogenesis and hydrogenotrophic pathway predominance

Across all wetland sites, we collected three intact soil cores in PVC tubes from 0 – 10 cm below the water table at the beginning, middle, and end of 35 m transects at each site. The samples were topped with site water and tightly sealed to ensure an anaerobic environment and transported to the University of Oregon. Anaerobic incubations took place at field temperatures (28 °C) within two days following arrival at University of Oregon. In a glove box filled with a N2 atmosphere (<5% H_2_ in the presence of palladium catalyst; Coy Laboratory Products, Grass Lake, Michigan, USA), approximately 10 g of wet weight soil were added to 120 mL serum bottles and mixed with 10 mL of deoxygenated, deionized water. Sample bottles were then flushed with N2 for 15 minutes to begin the incubation. Headspace samples were analyzed over the course of three days (0, 1, 2, and 3 days). Total CH_4_ was calculated using Henry’s Law, adjusting for solubility, temperature, and pH (Bridgham & Ye, 2013). CH_4_ production rates were calculated using the linear accumulation (r^2^ > 0.90 in all cases) of gas through time. To measure hydrogenotrophic methanogenesis, we used a ^14^CO_2_ tracer method (Keller & Bridgham, 2007), which allowed us to distinguish the percent of total CH_4_ that had been produced using CO_2_/H_2_ as the substrate (i.e. hydrogenotrophic methanogensis). This was done with a gas chromatograph fitted with a radioactive gas proportional counter (LabLogic Systems) simultaneously with the gross methanogenesis measurements over three days.

### Soil physical and chemical analysis

We measured a suite of abiotic variables from each soil sample to incorporate into our community model as a covariate (see below). For the methanogenesis study, we recorded pH, total % nitrogen (N), organic carbon (C), moisture content, and bulk density. For the methanotrophy study, we measured N, C, moisture content, and bulk density. Soil pH was determined from a 1:1 soil to deionized water solution. Total % N and organic C were measured on a Costech ECS 4010 Elemental Analyzer (Valencia, CA), with each sample analyzed in duplicate. Moisture content (gravimetric water content) and bulk density were measured by the change in weight of a soil sub-sample following 48 hours of drying at 60 °C. Additionally, material from the three soil cores at each site was combined and homogenized for texture analysis using the hydrometer method (Gavlak, Horneck, Miller, & Kotuby-Amarcher, 2003), with 5% sodium hexametaphosphate as the dispersing solution. Because soil texture measurements required pooling samples from each site- and all other variables were measured on a per sample basis, texture was not included in the environmental covariate calculation.

### Soil RNA/DNA co-extraction and sequencing

Soil DNA and RNA were co-extracted from Lifeguard-preserved soil samples using MoBio’s Powersoil RNA Isolation kit with the DNA Elution Accessory Kit (MoBio, California, USA) following manufacturer’s instructions. RNA was reverse transcribed to cDNA using Superscript III first-strand reverse transcriptase and random hexamer primers (Life Technologies, USA). Extractions were quantified using Qubit (Life Technologies, USA). We amplified the V4 region of the 16S SSU rRNA gene in sample DNA and cDNA using the primers 515F and 806R (Caporaso et al., 2011). Sequencing libraries were prepped using a dual-indexing approach (Fadrosh et al., 2014; Kozich, Westcott, Baxter, Highlander, & Schloss, 2013). In short, each PCR reaction was performed using 12.5 μl NEBNext Q5 Hot Start HiFi PCR master mix (New England Biolabs, USA), 11.5 μl gene-specific primer mix (1.09 μM of each primer), and 1 μl template (DNA or cDNA). A subset of samples was used to find 1) the optimal primer annealing temperature, and 2) the minimum number of cycles for adequate target amplification. For the 16S rRNA gene target this was 61° C and 20 cycles. The final reaction conditions were: 98° C 30 seconds (initialization), 98° C 10 seconds (denaturation), gene-specific annealing step for 20 seconds (see above), and 72° C for 20 seconds (final extension). Reactions were followed by magnetic bead purification using 20 μl Mag-Bind RxnPure Plus isolation beads (Omega Bio-Tek, USA). Reactions were quantified using Qubit (Life Technologies, USA) then were multiplexed at equimolar concentration. Final pooled amplicon libraries were sequenced using the Illumina MiSeq (300 paired-end) platform at the Oregon State University Center for Genome Research and Biocomputing facility.

### Bioinformatic processing

Demultiplexed reads were joined using PEAR (version 0.9.10) with default parameters (Zhang, Kobert, Flouri, & Stamatakis, 2014) and quality filtered using Prinseq (version 0.20.4) (Schmieder & Edwards, 2011). Sequences with a mean quality score ≥ 30 and length 250-350 bp were retained. Sequences were dereplicated, denoised, and checked for chimeras using the DADA2 pipeline (Version 1.6) (Callahan et al., 2016) implemented in QIIME2 (Caporaso et al., 2010) (https://qiime2.org). For the methanogenesis study 17 of the 18 DNA-based samples, and 13 of the 18 RNA-based samples had sufficient quality to be included. For the methanotrophy study 41 of the 44 DNA-based samples, and 34 of the 44 RNA-based samples passed our quality filter. Taxonomy was assigned to the resulting amplicon sequence variants (ASVs) using the Ribosomal Database Project online classifier (Wang, Garrity, Tiedje, & Cole, 2007) with database release 11.5 (Cole et al., 2014).

### Quantitative PCR

We quantified the abundance of methanogens in each sample from which CH_4_ production and hydrogenotrophic pathway predominance were measured. To do so we performed qPCR of the *mcrA* gene on sample DNA using the mlas-mcrA-rev primer combination (Steinberg & Regan, 2008). Samples were run on an ABI StepOnePlus thermocycler (ABI, USA), using Kapa SYBR reagents (Kapa Biosystems, USA) according to manufacturer recommendations. For each sample, 8 ng DNA was used, and the following amplification conditions were applied following optimization: 98° C 10 minutes, 98° C 15 seconds, 55.6° 15 seconds, 72° 60 seconds. A melt curve analysis was performed to verify target amplification. Not all samples reliably amplified; in the end we obtained useable *mcrA* quantifications for 12 of the 18 samples. Although attempted, the transcriptional activity of *mcrA* from soil cDNA did not produce reliable results, and thus is not included in this study. We used a similar approach to quantify methanotroph abundance and transcriptional activity by targeting the *pmoA* gene in the sample DNA and cDNA, respectively, using the A189 – mb661 primer combination (Bourne, Donald, & Murrell, 2001). Reactions were performed on a Bio-Rad CFX96 real-time qPCR instrument, using SsoAdvanced Universal SYBR Green supermix (Bio-Rad, USA) following manufacturer instructions. For each sample, 2 ng template were used with the following reaction conditions: 98° C 10 minutes, 98° C 15 seconds, 55.6° 15 seconds, 72° 60 seconds.

All samples were amplified in triplicate. In both cases, sample amplification was compared to a standard positive control to quantify total gene (or transcript) copy number. We obtained useable *pmoA* quantifications for 42 of the 44 DNA samples and 32 of the 44 RNA samples. In the case of *mcrA,* the positive control was an *mcrA* plasmid and, in the case of *pmoA,* we used purified DNA from strain *Methylococcus capsulatus* Foster and Davis (ATCC 33009D-5). We used LinRegPCR (Ramakers, Ruijter, Deprez, & Moorman, 2003; Ruijter et al., 2009) to process amplification data which allows for the calculation of individual PCR efficiencies. Individual PCR efficiencies did not significantly differ between habitat types, so gene copy was calculated using the average PCR efficiency of all reactions. Finally, gene copy (or transcript) numbers were normalized to the total ng DNA (or cDNA) used in the reaction.

### Microbial community attribute calculations, statistical analysis, and comprehensive modeling

All statistical analyses were performed using R (version 3.4.4) (R Core Team, 2018). Community matrices for the methanotrophy experiments were rarefied ten times at 28,000 counts per sample, then averaged, and community matrices for the methanogenesis experiments were rarefied to 30,400 counts per sample ten times and averaged, in order to account for differences in sampling extent across samples. Additionally, these rarefied community matrices were subsetted for known methanogens/methanotrophs (Supplementary Table 1) to create functional group community matrices. We categorized community data into the following four attributes (Table 1): 1) *Abundance* of methanogens/methanotrophs using qPCR of *mcrA* and *pmoA* genes (described above) respectively, as well as the relative abundance of known methanogens/methanotrophs in the DNA-inferred 16S-based prokaryotic community, 2) *Activity* of methanogens/methanotrophs using the relative abundance of methanogens and methanotrophs in the RNA-inferred prokaryotic community, and qPCR of *pmoA* using soil cDNA (described above), 3) *Composition* of the broader prokaryotic community or of subsets of known methanogen/methanotroph taxa from the 16S-inferred community matrix (described below), and 4) *Diversity* of the prokaryotic community as well as methanogen/methanotroph community subsets using ASV-level richness and Shannon diversity. Species richness and Shannon diversity were calculated using the ‘vegan’ package (Oksanen et al., 2015) in R. Regression plots were created using the package ‘ggplot2’ (Wickham, 2009), and 95% confidence interval bands were plotted around each linear fit using the ‘geom_smooth’ function (option = ‘lm’). Difference in process rates among wetlands and uplands was assessed using a Kruskal-Wallis test.

**Table 1:** The community attributes calculated in this study used to identify putative community controls on process rates.

| Attribute | Variables |
| --- | --- |
| Abundance | qPCR (of *mcrA* & *pmoA* genes)<br>Relative abundance (in DNA-inferred prokaryotic community) of methanogens/methanotrophs |
| Activity | cDNA qPCR (*pmoA*)<br>Relative abundance (in RNA-inferred prokaryotic community) of methanogens/methanotrophs |
| Composition | Matrix of 16S-based prokaryotic community inferred from RNA<br>Matrix of 16S-based prokaryotic community (DNA-inferred)<br>Matrix of methanogen/methanotroph taxa subsetted from 16S-based matrix (DNA-inferred)<br>Species (ASV) richness of RNA/DNA-inferred prokaryotic community |
| Diversity | ASV richness of methanogen/methanotroph community subsets (RNA & DNA)<br>Shannon (ASV) diversity of RNA/DNA-inferred prokaryotic community<br>Shannon diversity of methanogen/methanotroph community subsets (RNA & DNA) |

#### Associations between community attributes and CH_4_-cycling processes

Because communities tend to have complex covariance structure, we took several measures to avoid spurious associations when distinguishing biotic correlates of CH_4_-cycling processes. First, our process measurements were performed under constant temperatures similar to those experienced in the field (described above). By doing so, we minimized the influence of certain environmental factors (*e.g.* temperature fluctuations or precipitation events) on our process rate measurements. Secondly, we added two covariates to our model when assessing associations with process rate measurements: 1) a soil physicochemical covariate, and 2) and a sample (community) covariate. The soil physicochemical covariate was calculated using principle components analysis (PCA) on soil physicochemical measurements (described above). This was done using the prcomp function in the ‘stats’ package of R (R Core Team, 2018), with variables scaled to unit variance to account for differences in units of measurement. Sample scores for the first principle component (PC1) were used as the environmental covariate. We applied a similar approach to calculate the community covariate. In this case, the rarefied community matrix was Hellinger transformed, then ordinated using the prcomp function while scaling to unit variance, in order to calculate the community-based sample PC1 scores. By including these covariates in our model we are regressing each community attribute against the residual variation of the CH_4_-cycling process of interest after accounting for the variation explained by environment and underlying sample structure.

To assess associations between community membership and process rate variation we used the regression approach described above on the relative abundance of individual taxa (ASVs). Since performing this type of analysis on thousands of individual taxa increases the risk of false positives, we adjusted our significance threshold (alpha = 0.05) using a Bonferroni correction, i.e. by dividing by the number of of taxa being tested. In cases where significant taxa were identified, those taxa were subsetted from the rarefied community matrix, then ordinated (following the above procedure) to reduce them to a single variable. The sample scores of the first principle component were subsequently used for regression against CH_4_-cycling process rates. Linear regressions were performed using the lm function in the ‘stats’ package in R. Gross methanotrophy and methanogenesis rate data were log2-transformed in order to satisfy assumptions of normality. For each relationship we report the adjusted R^2^ and *p* value.

## RESULTS

### Identifying biotic drivers of CH_4_-cycling processes Methanogenesis processes

#### Gross Methanogenesis

Gross methanogenesis rates varied from 0.06 to 6.7 μmol CH_4_ g soil^-1^ d^-1^ (average 1.1 ± 1.5 μmol CH_4_ g soil^-1^ d^-1^, Fig. 1A), with rates in mineral soil wetlands on average 3.8 times higher than peatlands. For breakdown of soil physico-chemical variables by habitat see Supplementary Table 2. We identified two community attributes that were associated with gross methanogenesis rate after accounting for sample and environmental covariance structure: methanogen abundance (i.e. the abundance of *mcrA* genes, Adj. R^2^ = 0.47, *p* = 0.008, Fig. 1B, Table 2), and the alpha diversity of methanogens (Richness: Adj. R^2^ = 0.25, *p* = 0.023, Shannon: Adj. R^2^ = 0.23, *p* = 0.029). Both relationships were positive, suggesting that more abundant and/or diverse populations of methanogens are associated with higher rates of gross methanogenesis. We did not identify any individual prokaryotic taxa that were significantly associated with gross methanogenesis after accounting for environmental and sample covariance and adjusting for multiple comparisons. Soil pH was the only significant abiotic predictor of methanogenesis (Adj. R^2^ = 0.24, *p* = 0.023), showing a positive relationship whereby values approaching neutral pH had higher rates of methanogenesis.

**Fig. 1:**
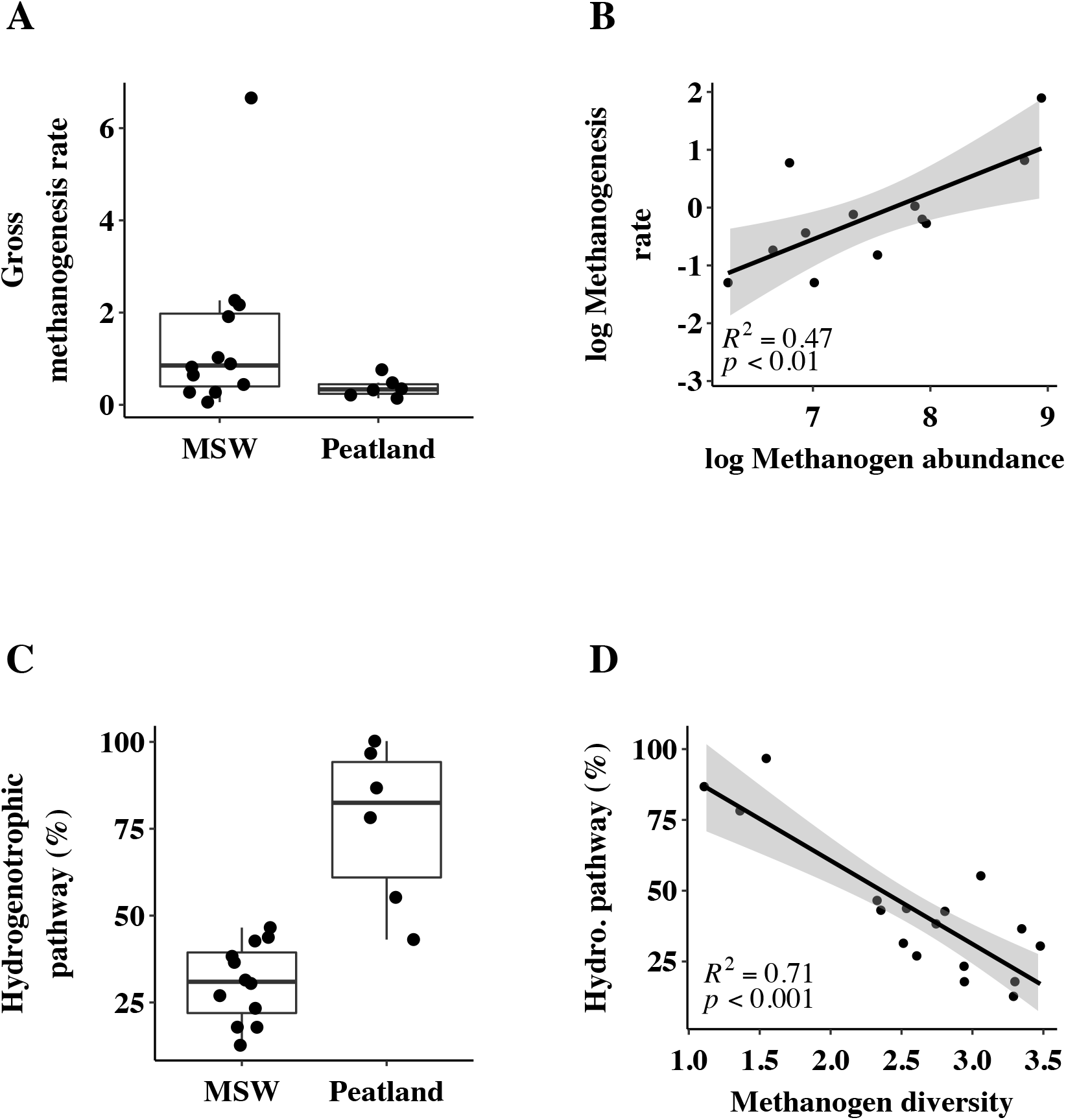
The rates and biotic drivers of methanogenesis-related processes across Congo Basin wetlands. A) Methanogenesis rate (μmol CH_4_ g soil^-1^ d^-1^) across the two wetland habitats. MSW = mineral soil wetland. B) The log_2_ methanogenesis rate is best predicted by log_2_ methanogen abundance (qPCR of *mcrA* gene). C) The predominance (%) of the hydrogenotrophic pathway in gross methanogenesis across wetland habitats. D) Hydrogenotrophic pathway predominance is inversely related to the alpha diversity (Shannon index) of the methanogen community. Each point represents an individual soil sample. Gray bands represent 95% confidence intervals of the linear model.

**Table 2:** Narrow scale processes have a higher association with community attributes. Process assessment scale: whether a process assessment is the aggregate result of several processes (broad) or only one (fine). Adj. R^2^ = the proportion of the variance explained by the attribute.

|  | Physiological<br>scale of<br>process | Best fit<br>community<br>attribute | Adj. R<sup>2</sup> | P value |
| --- | --- | --- | --- | --- |
| ***Methanotrophy processes*** |  |  |  |  |
| Gross Methanotrophy<br>(V<sub>max</sub>) | Broad | Abundance<br>(*pmoA* copy) | 0.14 | 0.01 |
| High affinity methanotrophy<br>(k) | Fine | Composition<br>(RNA-inferred) | 0.64 | < 0.001 |
| ***Methanogenesis processes*** |  |  |  |  |
| Gross Methanogenesis<br>(μmol CH<sub>4</sub> g soil<sup>-1</sup> d<sup>-1</sup>) | Broad | Abundance<br>(*mcrA* copy) | 0.47 | 0.008 |
| Hydrogenotrophic<br>methanogenesis<br>(% gross CH<sub>4</sub> production) | Fine | Diversity<br>(methanogens) | 0.79 | < 0.001 |

#### Hydrogenotrophic methanogenesis

The most pronounced difference in the predominance of the hydrogenotrophic pathway was between mineral soil wetlands and peatlands, where it accounted for 30.7 ± 11.2 % and 76.7 ± 23.0 % of total methanogenesis, respectively (Fig. 1C). We identified a negative relationship between the DNA-inferred alpha diversity of the methanogens and the predominance of the hydrogenotrophic pathway (Shannon: Adj. R^2^ = 0.710, *p* < 0.001 Fig. 1D, Table 2, Richness: Adj. R^2^ = 0.455, *p* = 0.002). We also identified a negative relationship with the RNA-inferred alpha diversity of the prokaryotic community (Shannon: Adj. R^2^ = 0.609, *p* = 0.001). The only abiotic variables that were significantly associated with hydrogenotrophic methanogenesis were a negative relationship with soil pH (Adj. R^2^=0.32, *p* < 0.01), and a positive relationship with soil C (Adj. R^2^=0.341, *p* < 0.01).

### Methanotrophy processes

#### Gross methanotrophy

Average rates of gross CH_4_ consumption (V_max_) were higher in wetland sites (including seasonal wetlands) (0.3 ± 0.05 μmol CH_4_ g soil^-1^ d^-1^) than upland sites (0.1 ± 0.02 μmol CH_4_ g soil^-1^ d^-1^) (Fig. 2A, Kruskal-Wallis: Chi-squared = 6.85, df = 1, *p* = 0.008), and significantly correlated with every abiotic variable measured. Gross methanotrophy rates were negatively correlated with soil bulk density (Adj. R^2^= 0.135, *p* =0.008). Note that the inverse of V_max_ is presented, i.e. higher uptake rates are represented by higher values. Conversely gross methanotrophy rates were positively correlated with soil moisture (Adj. R^2^= 0.22, *p* < 0.001), soil C (Adj. R^2^= 0.094, *p* =0.024), and soil N (Adj. R^2^= 0.105, *p* =0.02). The only community attribute that correlated with gross methanotrophy after accounting for variation due to soil physicochemical variables was methanotroph abundance (i.e. *pmoA* copy number, Adj. R^2^=0.139, p=0.01, Fig. 2B, Table 2).

**Fig. 2.**
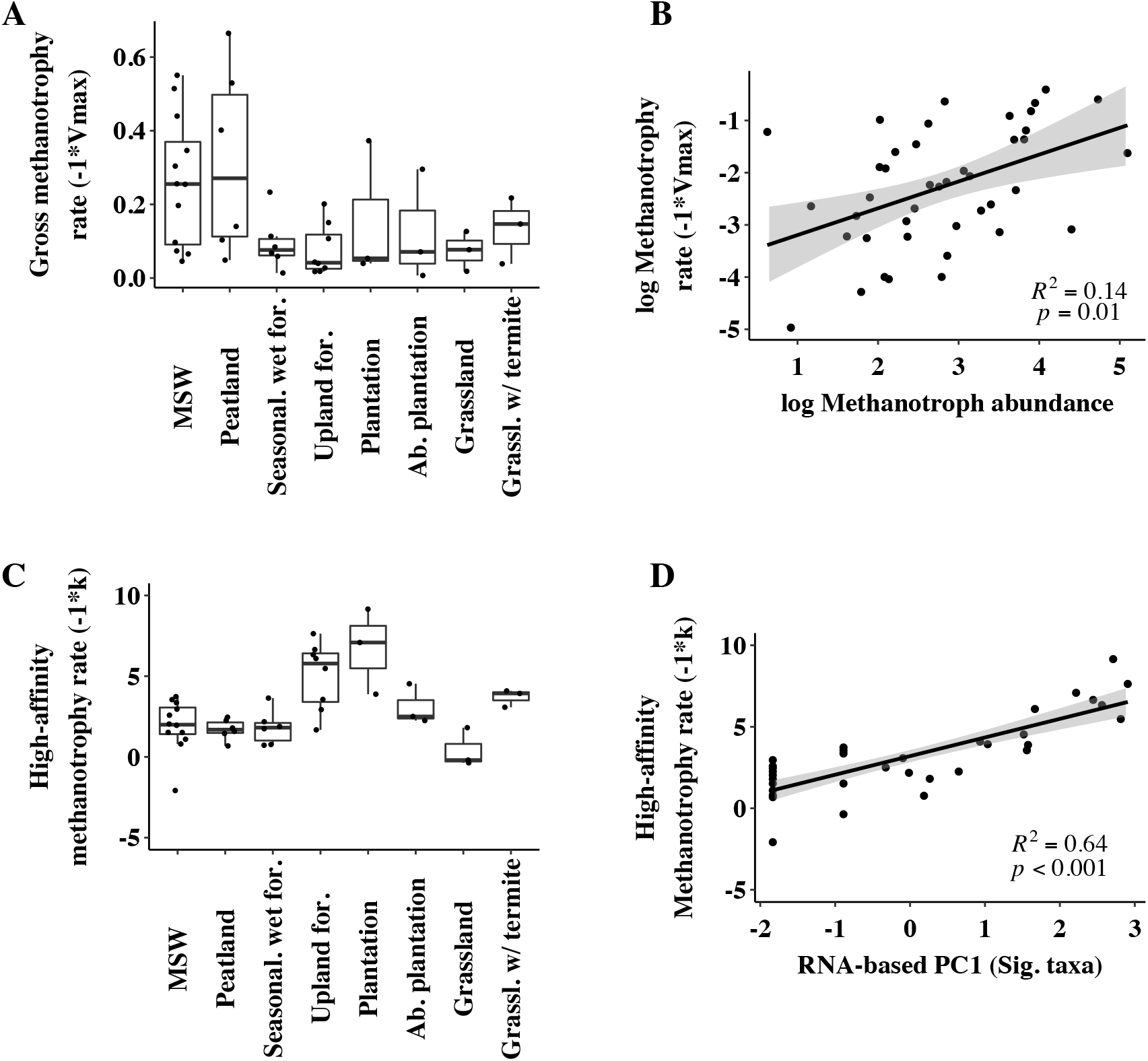
The rates and biotic drivers of methanotrophy-related processes across Congo Basin habitats. Note the inverse of V_max_ and k and presented so that higher positive values correspond to higher rates of uptake. A) Gross methanotrophy rate (−1* V_max_) across habitats. MSW = mineral soil wetland, Seasonal. Wet for. = seasonally inundated forest, Upland for. = upland forest, Ab. plantation = abandoned plantation, Grassland w/ termite = grassland habitat dominated by termite mounds. B) Gross methanotrophy rate is positively related to the log_2_ abundance of methanotrophs (*pmoA* gene copy number). C) High-affinity methanotrophy rate across habitats. D) High-affinity methanotrophy rates are best predicted by the first principle component (PC1) of a subset of significantly associated taxa (Sig. taxa) identified in the RNA-inferred prokaryotic community. Gray bands represent 95% confidence intervals of the linear model.

#### High-affinity methanotrophy

Of the four processes measured, high-affinity methanotrophy was the only process to exhibit composition-based control involving the broader prokaryotic community beyond known methanotrophs. Rates of high-affinity methane oxidation varied considerably across our samples (range −9.15 to 2.08) (Fig. 2C) and were higher in upland sites relative to wetland sites (including seasonal wetlands) (Kruskal-Wallis: Chi-squared = 11.52, df = 1, *p* < 0.001). After accounting for sample and environmental covariance, we identified 11 taxa in the DNA-inferred community and 5 taxa in the RNA-inferred community whose relative abundance significantly corresponded to high-affinity methanotrophy rates. The finest level of taxonomic assignment for these taxa varied from order to genus. DNA-inferred taxa identified included members of the genera *Nitrosphaera* (Thaumarchaeota), *Roseiarcus* (Alphaproteobacteria), *Aminiphilus* (Synergistetes), and *Acidisoma* (Alphaproteobacteria), a member of the family Thermosporaceae (Actinobacteria), four members of Acidobacteria Group 2, and two members of Acidobacteria Group 6 (Supplementary Table 3). There was no ASV-level overlap with the RNA-inferred taxa that were identified. The RNA-inferred taxa included members of the genera *Beijerinckia* (Alphaproteobacteria), *Deferrisoma* (Deltaproteobacteria), *Nitrospirillum* (Alphaproteobacteria), the family Thermonosporaceae (Actinobacteria), and a member of Acidobacteria Group 1 (Supplementary Table 3).

We next ordinated the relative abundances of these taxa to reduce them to a single variable. The first principle component of both of these community subsets significantly correlated with high-affinity methanotrophy (RNA-inferred PC1 axis: Adj. R^2^ = 0.647, *p* < 0.001, Fig. 2D, Table 2, DNA-inferred PC1 axis: Adj. R^2^ = 0.583, *p* < 0.001). Lastly, we tested if high-affinity methanotrophy correlated with any other community attributes. After accounting for covariance structure, there were no other biotic correlates of high-affinity methanotrophy beyond composition-based variables.

## DISCUSSION

### The utility of microbial community attributes in predicting function

The utility of conventional microbial community measurements for predicting ecosystem functions has been recently called into question (Graham et al., 2014; Inkpen et al., 2017; Louca et al., 2018; Rocca et al., 2014). Here we argue that many of the difficulties associated with detecting community structure – ecosystem function relationships could be overcome with careful consideration of the community attributes chosen, the physiological scale of functional assessment, and how environmental variation is controlled. In this study we have addressed each of these concerns. Our work demonstrates 1) that there is a connection between communities and ecosystem processes after accounting for variation due to external factors, and 2) that the strength, as well as the nature, of the relationship between communities and ecosystem function depends on the physiological scale of the function.

Ecosystem function measurements face issues of physiological scale (Inkpen et al., 2017), i.e. they represent the aggregate result of numerous underlying processes. Detecting a causal link between community measurements and ecosystem function thus becomes increasingly difficult for functions that include inputs from multiple processes. This suggests that as one narrows the functional scale to a level that more closely approximates individual physiologies, the strength of the relationship with the community should increase. Our study supports this hypothesis. For both methanogenesis- and methanotrophy-related processes, a similar trend emerged; broader-scale functions had a weaker relationship with community attributes than finer-scale functions. For instance, the best predictor of gross methanogenesis was methanogen abundance (i.e. *mcrA* copy number), explaining 47% of the variance, whereas hydrogenotrophic methanogensis – which represents a finer physiological scale-showed the strongest association with methanogen alpha diversity, explaining 71% of the variance (Table 2). For methanotrophy processes, gross methanotrophy was best predicted by methanotroph abundance (i.e. *pmoA* copy number), but this connection was quite weak and only explained ~14% of the variance. By contrast, much more of the variance in high-affinity methanotrophy could be explained by RNA-inferred community composition (~64%, Table 2). While our specific hypothesis only represents two processes for each type of function, our results provide support for the idea that the physiological scale of function is an important consideration for community structure-ecosystem function studies. Future work could extend the generality of our findings by assessing an even wider assortment of physiological scales (e.g. net CH_4_ flux) or by extending this framework to other ecosystem functions (e.g. soil respiration).

### Developing community variables that represent putative process controls

Communities can control processes in several distinct ways. It is therefore important that the variables we choose to represent communities reflect these types of control. One key finding from our work was that broader scale processes had distinctly different predictors than the underlying finer scale processes. Hence, several of the relationships uncovered by our work would have gone undetected had we focused solely on a single attribute. Both gross methanogenesis and methanotrophy were best predicted by abundance-based variables (i.e. *mcrA* and *pmoA* copy number, respectively), suggesting that these processes could be limited by the total number of individuals capable of performing them. Similar relationships between abundance and methanotrophy/methanogenesis rates have been reported (Freitag & Prosser, 2009; Ma et al., 2012), although a number of studies have demonstrated no such relationship (see Rocca et al., 2014) – suggesting other important controls. Hydrogenotrophic methanogenesis and high-affinity methanotrophy – the finer-scale processes – by contrast were best predicted by diversity and composition measures, respectively. Hydrogenotrophic predominance was inversely related to methanogen diversity. One possible explanation for this relationship is that more diverse methanogen consortia would be more likely to contain other methanogen functional groups (e.g. acetoclastic or methylotrophic taxa), which would thus affect the relative contribution of the hydrogenotrophic pathway to gross methanogenesis.

The strongest – and only – biotic variables associated with high-affinity methanotrophy were based on RNA- and DNA-inferred composition (i.e. taxon identity), suggesting that trait differences among taxa could be especially important for this process. Interestingly, none of the associated taxa we identified were known methanotrophs. This insight highlights the importance of not restricting our search to taxa that are canonically associated with methanotrophy. While there are issues with the commensurability of taxonomic composition and ecosystem function (Inkpen et al., 2017), these taxa could still be causally linked to high-affinity methanotrophy in at least two ways. First, at least some could be non-canonical methanotrophs that are consuming CH_4_. Even though the ability to consume CH_4_ is conserved phylogenetically (Martiny et al., 2013), canonical methanotrophs (as well as high affinity methanotrophs) are nevertheless polyphyletic (Knief, 2015), suggesting that methanotrophy has evolved multiple times or has been laterally transferred. Another possibility is that these taxa influence methanotrophy through ecological interactions with methanotrophs (e.g. competition or facilitation). A final possibility is that these taxa are simply a marker for high-affinity methanotrophy, co-varying with a variable that is causally linked. One of the DNA-inferred taxa identified, *Nitrosphaera sp.* (Thaumarchaeota), is a known ammonia oxidizer. Ammonia monooxygenase and methane monooxygenase are phylogenetically related (Holmes, Costello, Lidstrom, & Murrell, 1995), and it has been suggested that ammonia oxidizers may be able to bind CH_4_ (Arp, Sayavedra-Soto, & Hommes, 2002; Ross & Rosenzweig, 2017). Another taxon, *Roseiarcus* (Alphaproteobacteria), is closely related to alphaproteobacterial methanotrophs and has been isolated from a methanotrophic consortium enriched from *Sphagnum* peat (Kulichevskaya, Danilova, Tereshina, Kevbrin, & Dedysh, 2014). Thus, *Roseiarcus* could be either directly consuming CH_4_, or interacting with methanotrophs. Two of the RNA-inferred taxa, *Nitrospirillum* and *Beijerinckia* (both Alphaproteobacteria), are diazotrophs and could thus influence methanotroph activity by fixing atmospheric N2. The influence of other correlated taxa on methanotrophy is less clear, but competition for soil nutrients/O_2_, facilitation, or simply acting as a marker remain possibilities.

While we attempted to equally represent each attribute in our modeling, activity-based attributes were under-represented in this study. Transcriptional levels of *pmoA* and *mcrA* have been used as proxies for methanotroph/methanogen activity in a variety of studies (Chen, Dumont, Cébron, & Murrell, 2007; Freitag & Prosser, 2009; Freitag et al., 2010). While we successfully quantified *pmoA* transcriptional activity, our attempt to do so for *mcrA* was problematic. This was not likely due to a lack of activity of methanogens (since CH_4_ was continuously produced throughout the experiment), but rather due to PCR inhibitors such as humic substances present in organic-rich soils. We took our community inference methods a step further than many other studies by using both RNA- and DNA-based 16S rRNA profiling. It has been established that the relative abundances of taxa in the RNA-inferred community should not be used as a direct assessment of activity levels (Papp, Mau, et al., 2018; Papp, Hungate, et al., 2018). However, it is likely that the RNA-inferred community is at least enriched for the active fraction of soil taxa relative to the DNA-inferred community (which could contain higher proportions of inactive, dormant, or dead individuals) (Lennon & Jones, 2011). Thus, by including RNA-based community inference we were able to use an additional window onto the community that could have functional relevance.

### Gaining a better understanding of tropical CH_4_ dynamics

Despite the established importance of tropical ecosystems as global CH_4_ sources (Kirschke et al., 2013; Sjögersten et al., 2014), our understanding of the ecological controls over CH_4_-cycling throughout these regions remains limited (Bridgham et al., 2013). The majority of studies aimed at identifying controls on CH_4_-cycling dynamics have been performed in higher latitude ecosystems such as northern peatlands (Bridgham et al., 2013). The contribution of biotic variables to CH_4_-cycling dynamics may be a more important determinant of variation across tropical environments relative to higher latitudes. Methanogen population growth rates are slow because all methanogenic pathways have very low thermodynamic yields that are often barely above the threshold for growth under *in-situ* conditions (Conrad, 1999; Megonigal, Mines, & Visscher, 2004). Thus, methanogens inhabiting high latitude environments must recover each season from freeze-thaw events and will exhibit low levels of activity/growth until warmer temperatures are reached. In tropical ecosystems where prolonged freezing conditions are essentially absent, trait-level distributions of CH_4_-cycling taxa could play a stronger role regulating spatial differences in CH_4_ dynamics. While our study was not designed to compare differences between high- and low-latitude ecosystems, an interesting trend that emerged from our results was that abiotic variables tended to be less predictive of CH_4_ cycling relative to biotic variables (with the exception of gross methanotrophy). Importantly, our laboratory incubations deliberately downplayed the influence of several key environmental factors (e.g. temperature fluctuations or rainfall events) that could influence CH_4_-cycling rates. Nevertheless, most of our environmental variables were independent from our incubation conditions (e.g. %C, %N, pH, soil bulk density), and each only explained a relatively small fraction of the variance. Thus, an important outstanding question is whether the strength and nature of biotic controls on CH_4_-cycling dynamics differs between high- and low-latitude ecosystems.

### Conclusion

How communities map onto ecosystem function is a widely debated topic. Central to this debate are issues related to the physiological scale of function, the identification of useful community attributes, and accounting for the underlying influence of the environment. Our study provides two unique perspectives to this debate. First, we suggest that broad definitions of function could be obscuring the connections with community attributes. Secondly, by compiling community attributes according to their putative role in controlling processes, we show that finer scale processes may face controls that differ from those of the coarser processes to which they contribute. Collectively, these findings demonstrate the importance of carefully defining the physiological scale of ecosystem function and performing community measurements that represent putative controls on ecosystem processes.

## ACKNOWLEDGEMENTS

We thank the Government of Gabon, Centre National de la Recherche Scientifique et Technologique for permission (Permit No AR0035/14/MESRS/CENAREST/CG/CST/CSAR) to conduct this study. We thank F. Bivigou, H. Memiaghe, L. Tchignoumba, E. Tobi, I. Akendengue for their help collecting samples. The University of Oregon, the Gabon-Oregon Transnational Center on Environment and Development, the Smithsonian Conservation Biology Institute, and Shell Gabon provided financial and logistical support. BB and KM are grateful for support by the National Science Foundation – Dimensions of Biodiversity program (DEB 14422214), and SB and AH are grateful for support by the Office of Biological and Environmental Research, Terrestrial Ecosystem Science Program, U.S. Department of Energy (DE-SC0008092 and DE-39SC0012088).

## DATA ACCESSIBILITY

Sequence data, sample meta data, biogeochemical data, ASV community matrices and R script for analysis will be made available upon acceptance of publication.

## AUTHOR CONTRIBUTIONS

KM, AH, SB and BB designed and performed research. KM, AH, AK, and AM analyzed data. KM wrote the paper with contributions from all authors.

## REFERENCES

Arp, D. J., Sayavedra-Soto, L. A., & Hommes, N. G. (2002). Molecular biology and biochemistry of ammonia oxidation by Nitrosomonas europaea. Archives of Microbiology, 178(4), 250–255. http://doi.org/10.1007/s00203-002-0452-0

Baldrian, P., Kolařík, M., Stursová, M., Kopecký, J., Valášková, V., Větrovský, T., … Voříšková, J. (2012). Active and total microbial communities in forest soil are largely different and highly stratified during decomposition. The ISMEJournal, 6(2), 248–58. http://doi.org/10.1038/ismej.2011.95

Bannar-Martin, K. H., Kremer, C. T., Ernest, S. M., Leibold, M. A., Auge, H., Chase, J., … Supp, S. R. (2018). Integrating community assembly and biodiversity to better understand ecosystem function: the Community Assembly and the Functioning of Ecosystems (CAFE) approach. Ecology Letters, 21, 167–180. http://doi.org/10.1111/ele.12895

Bier, R. L., Bernhardt, E. S., Boot, C. M., Graham, E. B., Hall, E. K., Lennon, J. T., … Wallenstein, M. D. (2015). Linking microbial community structure and microbial processes: An empirical and conceptual overview. FEMS Microbiology Ecology, 91(10), 1–11. http://doi.org/10.1093/femsec/fiv113

Bodelier, P. L. E., Meima-Franke, M., Hordijk, C. a, Steenbergh, A. K., Hefting, M. M., Bodrossy, L., … Seifert, J. (2013). Microbial minorities modulate methane consumption through niche partitioning. The ISME Journal, 7(11), 2214–28. http://doi.org/10.1038/ismej.2013.99

Bodelier, P. L. E., Roslev, P., Henckel, T., & Frenzel, P. (2000). Stimulation by ammonium-based fertilizers of methane oxidation in soil around rice roots. Nature, 403, 421–424.

Bourne, D. G., Donald, I. A. N. R. M. C., & Murrell, J. C. (2001). Comparison of pmoA PCR Primer Sets as Tools for Investigating Methanotroph Diversity in Three Danish Soils. Applied and Environmental Microbiology, 67(9), 3802–3809. http://doi.org/10.1128/AEM.67.9.3802

Bridgham, S. D., Cadillo-Quiroz, H., Keller, J. K., & Zhuang, Q. (2013). Methane emissions from wetlands: Biogeochemical, microbial, and modeling perspectives from local to global scales. Global Change Biology, 19(5), 1325–1346. http://doi.org/10.1111/gcb.12131

Bridgham, S. D., & Ye, R. (2013). Organic Matter Mineralization and Decomposition. In R. D. DeLaune, K. R. Reddy, C. J. Richardson, & J. P. Megonigal (Eds.), Methods in Biogeochemistry of Wetlands (pp. 385–406). Madison, WI: SSSA.

Callahan, B. J., McMurdie, P. J., Rosen, M. J., Han, A. W., Johnson, A. J. A., & Holmes, S. P. (2016). DADA2: High-resolution sample inference from Illumina amplicon data. Nature Methods, 13, 581. Retrieved from http://dx.doi.org/10.1038/nmeth.3869

Caporaso, J. G., Kuczynski, J., Stombaugh, J., Bittinger, K., Bushman, F. D., Costello, E. K., … Knight, R. (2010). QIIME allows analysis of high-throughput community sequencing data Intensity normalization improves color calling in SOLiD sequencing. Nature Methods, 7(5), 335–336. http://doi.org/10.1038/NMETH.F.303

Caporaso, J. G., Lauber, C. L., Walters, W. A., Berg-lyons, D., Lozupone, C. A., Turnbaugh, P. J., … Knight, R. (2011). Global patterns of 16S rRNA diversity at a depth of millions of sequences per sample. Proceedings of the National Academy of Sciences, 108, 4516–4522. http://doi.org/10.1073/pnas.1000080107/-/DCSupplemental.www.pnas.org/cgi/doi/10.1073/pnas.1000080107

Chen, Y., Dumont, M. G., Cébron, A., & Murrell, J. C. (2007). Identification of active methanotrophs in a landfill cover soil through detection of expression of 16S rRNA and functional genes. Environmental Microbiology, 9(11), 2855–69. http://doi.org/10.1111/j.1462-2920.2007.01401.x

Cole, J. R., Wang, Q., Fish, J. A., Chai, B., McGarrell, D. M., Sun, Y., … Tiedje, J. M. (2014). Ribosomal Database Project: data and tools for high throughput rRNA analysis. Nucleic Acids Research, 42(D1), D633–D642. http://doi.org/10.1093/nar/gkt1244

Conrad, R. (1999). Contribution of hydrogen to methane production and control of hydrogen concentrations in methanogenic soils and sediments. FEMS Microbiology Ecology, 28(3), 193–202. http://doi.org/10.1016/S0168-6496(98)00086-5

Delmas, R. A., Tathy, J. P., & Cros, B. (1992). Atmospheric methane budget in Africa. Journal of Atmospheric Chemistry, 14(1), 395–409. http://doi.org/10.1007/BF00115247

Delmas, R., Servant, J., Tathy, J., Cros, B., & Labat, M. (1992). Sources and sinks of methane and carbone dioxide exchanges in mountain forest in Equatorial Africa. Journal of Geophysical Research, 97(90), 6169–6179.

Fadrosh, D. W., Ma, B., Gajer, P., Sengamalay, N., Ott, S., Brotman, R. M., & Ravel, J. (2014). An improved dual-indexing approach for multiplexed 16S rRNA gene sequencing on the Illumina MiSeq platform. Microbiome, 2(6), 1–7.

Freitag, T. E., & Prosser, J. I. (2009). Correlation of methane production and functional gene transcriptional activity in a peat soil. Applied and Environmental Microbiology, 75(21), 6679–87. http://doi.org/10.1128/AEM.01021-09

Freitag, T. E., Toet, S., Ineson, P., & Prosser, J. I. (2010). Links between methane flux and transcriptional activities of methanogens and methane oxidizers in a blanket peat bog. FEMS Microbiology Ecology, 73, 157–165. http://doi.org/10.1111/j.1574-6941.2010.00871.x

Garcia, J. L., Patel, B. K., & Ollivier, B. (2000). Taxonomic, phylogenetic, and ecological diversity of methanogenic Archaea. Anaerobe, 6(4), 205–26. http://doi.org/10.1006/anae.2000.0345

Gavlak, R., Horneck, D., Miller, R. O., & Kotuby-Amarcher, J. (2003). Soil, Plant, and Water Reference Methods for the Western Region. Western Region Extension Publication, WREP–125.

Graham, E. B., Wieder, W. R., Leff, J. W., Weintraub, S. R., Townsend, A. R., Cleveland, C. C., … Nemergut, D. R. (2014). Do we need to understand microbial communities to predict ecosystem function ? A comparison of statistical models of nitrogen cycling processes. Soil Biology and Biochemistry, 68, 279–282. http://doi.org/10.1016/j.soilbio.2013.08.023

Ho, A., Kerckhof, F.-M., Luke, C., Reim, A., Krause, S., Boon, N., & Bodelier, P. L. E. (2013). Conceptualizing functional traits and ecological characteristics of methane-oxidizing bacteria as life strategies. Environmental Microbiology Reports, 5(3), 335–45. http://doi.org/10.1111/j.1758-2229.2012.00370.x

Holmes, A. J., Costello, A., Lidstrom, M. E., & Murrell, J. C. (1995). Evidence that particulate methane monooxygenase and ammonia monooxygenase may be evolutionarily related. FEMS Microbiology Letters, 132, 203–208.

Inkpen, S. A., Douglas, G. M., Brunet, T. D. P., Leuschen, K., Doolittle, W. F., & Langille, M. G. I. (2017). The coupling of taxonomy and function in microbiomes. Biology & Philosophy, http://doi.org/10.1007/s10539-017-9602-2

Jones, S. E., & Lennon, J. T. (2010). Dormancy contributes to the maintenance of microbial diversity. Proceedings of the National Academy of Sciences of the United States of America, 107(13), 5881–6. http://doi.org/10.1073/pnas.0912765107

Kamke, J., Taylor, M. W., & Schmitt, S. (2010). Activity profiles for marine sponge-associated bacteria obtained by 16S rRNA vs 16S rRNA gene comparisons. The ISME Journal, 4(4), 498–508. http://doi.org/10.1038/ismej.2009.143

Keller, J., & Bridgham, S. (2007). Pathways of anaerobic carbon cycling across an ombrotrophic-minerotrophic peatland gradient. Limnology and Oceanography, 52(1), 96–107. http://doi.org/10.4319/lo.2007.52.1.0096

Kirschke, S., Bousquet, P., Ciais, P., Saunois, M., Canadell, J. G., Dlugokencky, E. J., … Zeng, G. (2013). Three decades of global methane sources and sinks. Nature Geoscience, 6(10), 813–823. Retrieved from http://dx.doi.org/10.1038/ngeo1955

Knief, C. (2015). Diversity and habitat preferences of cultivated and uncultivated aerobic methanotrophic bacteria evaluated based on pmoA as molecular marker. Frontiers in Microbiology, 6, 1–38. http://doi.org/10.3389/fmicb.2015.01346

Kolb, S. (2009). The quest for atmospheric methane oxidizers in forest soils. Environmental Microbiology Reports, 1(5), 336–346. http://doi.org/10.1111/j.1758-2229.2009.00047.x

Kozich, J. J., Westcott, S. L., Baxter, N. T., Highlander, S. K., & Schloss, P. D. (2013). Development of a dual-index sequencing strategy and curation pipeline for analyzing amplicon sequence data on the MiSeq Illumina sequencing platform. Applied and Environmental Microbiology, 79(17), 5112–20. http://doi.org/10.1128/AEM.01043-13

Kulichevskaya, I. S., Danilova, O. V, Tereshina, V. M., Kevbrin, V. V, & Dedysh, S. N. (2014). Descriptions of Roseiarcus fermentans gen. nov., sp. nov., a bacteriochlorophyll a-containing fermentative bacterium related phylogenetically to alphaproteobacterial methanotrophs, and of the family Roseiarcaceae fam. nov. International Journal of Systematic and Evolutionary Microbiology, 64(8), 2558–2565. https://ijs.microbiologyresearch.org/content/journal/ijsem/10.1099/ijs.0.064576-0

Lee, M. E., Alonso, A., Dallmeier, F., Campbell, P., & Pauwels, O. S. G. (2006). The Gamba Complex of Protected Areas : An Illustration of Gabon’s Biodiversity. Bulletin of the Biological Society of Washington, (12), 229–242.

Legendre, P. (1993). Spatial autocorrelation: trouble or new paradigm? Ecology, 74(6), 1659–1673.

Lennon, J. T., & Jones, S. E. (2011). Microbial seed banks: the ecological and evolutionary implications of dormancy. Nature Reviews Microbiology, 9(2), 119–30. http://doi.org/10.1038/nrmicro2504

Loreau, M., Naeem, S., Inchausti, P., Bengtsson, J., Grime, J. P., Hector, A., … Wardle, D. A. (2001). Biodiversity and Ecosystem Functioning: Current Knowledge and Future Challenges. Science, 294(5543), 804LP–808. Retrieved from http://science.sciencemag.org/content/294/5543/804.abstract

Louca, S., Polz, M. F., Mazel, F., Albright, M. B. N., Huber, J. A., Connor, M. I. O., … Parfrey, L. W. (2018). Function and functional redundancy in microbial systems. Nature Ecology & Evolution. http://doi.org/10.1038/s41559-018-0519-1

Ma, K., Conrad, R., & Lu, Y. (2012). Responses of methanogen mcrA genes and their transcripts to an alternate dry/wet cycle of paddy field soil. Applied and Environmental Microbiology, 78(2), 445–54. http://doi.org/10.1128/AEM.06934-11

MacDonald, J. A., Jeeva, D., Eggleton, P., Davies, R., Bignell, D. E., Fowler, D., … Maryati, M. (1999). The effect of termite biomass and anthropogenic disturbance on the CH4 budgets of tropical forests in Cameroon and Borneo. Global Change Biology, 5(8), 869–879.

Martiny, A. C., Treseder, K., & Pusch, G. (2013). Phylogenetic conservatism of functional traits in microorganisms. The ISME Journal, 7(4), 830–8. http://doi.org/10.1038/ismej.2012.160

McCalley, C. K., Woodcroft, B. J., Hodgkins, S. B., Wehr, R. a., Kim, E.-H., Mondav, R., … Saleska, S. R. (2014). Methane dynamics regulated by microbial community response to permafrost thaw. Nature, 514(7523), 478–481. http://doi.org/10.1038/nature13798

Megonigal, J. P., Mines, M. E., & Visscher, P. T. (2004). Anaerobic Metabolism: Linkages to Trace Gases and Aerobic Processes. In Biogeochemistry (Vol. 8, pp. 350–362). Gulf Professional Publishing.

Nazaries, L., Murrell, J. C., Millard, P., Baggs, L., & Singh, B. K. (2013). Methane, microbes and models: fundamental understanding of the soil methane cycle for future predictions. Environmental Microbiology, 15(9), 2395–417. http://doi.org/10.1111/1462-2920.12149

Nazaries, L., Pan, Y., Bodrossy, L., Baggs, E. M., Millard, P., Murrell, J. C., & Singh, B. K. (2013). Evidence of microbial regulation of biogeochemical cycles from a study on methane flux and land use change. Applied and Environmental Microbiology, 79(13), 4031–40. http://doi.org/10.1128/AEM.00095-13

Oksanen, J., Blanchet, F. G., Roeland, K., Legendre, P., Minchin, P., O’Hara, R. B., … Wagner, H. (2015). vegan: Community ecology package. Retrieved from http://cran.r-project.org

Papp, K., Hungate, B. A., & Schwartz, E. (2018). Microbial rRNA Synthesis and Growth Compared through Quantitative Stable Isotope Probing with H218O. Applied and Environmental Microbiology, 84(8), 1–11.

Papp, K., Mau, R. L., Hayer, M., Koch, B. J., Hungate, B. A., & Schwartz, E. (2018). Quantitative stable isotope probing with H218O reveals that most bacterial taxa in soil synthesize new ribosomal RNA. The ISME Journal, 18–20. http://doi.org/10.1038/s41396-018-0233-7

R Core Team. (2018). R: A language and environment for statistical computing. Vienna, Austria: R Foundation for statistical computing. Retrieved from http://cran.r-project.org

Ramakers, C., Ruijter, J. M., Deprez, R. H. L., & Moorman, A. F.. (2003). Assumption-free analysis of quantitative real-time polymerase chain reaction (PCR) data. Neuroscience Letters, 339(1), 62–66. http://doi.org/10.1016/S0304-3940(02)01423-4

Rocca, J. D., Hall, E. K., Lennon, J. T., Evans, S. E., Waldrop, M. P., Cotner, J. B., … Wallenstein, M. D. (2014). Relationships between protein-encoding gene abundance and corresponding process are commonly assumed yet rarely observed. The ISME Journal, 9(8), 1693–1699. http://doi.org/10.1038/ismej.2014.252

Ross, M. O., & Rosenzweig, A. C. (2017). A tale of two methane monooxygenases. Journal of Biological Inorganic Chemistry, 22(2-3), 307–319. http://doi.org/10.1007/s00775-016-1419-y

Ruijter, J. M., Ramakers, C., Hoogaars, W. M. H., Karlen, Y., Bakker, O., van den Hoff, M. J. B., & Moorman, a F. M. (2009). Amplification efficiency: linking baseline and bias in the analysis of quantitative PCR data. Nucleic Acids Research, 37(6), e45. http://doi.org/10.1093/nar/gkp045

Schimel, J. P. (1995). Ecosystem Consequences of Microbial Diversity and Community Structure. In F. S. Chapin III & C. Körner (Eds.), Arctic and Alpine Biodiversity: Patterns, Causes and Ecosystem Consequences (pp. 239–254). Berlin: Springer-Verlag.

Schimel, J. P., & Gulledge, J. (1998). Microbial community structure and global trace gases. Global Change Biology, 4, 745–758.

Schmieder, R., & Edwards, R. (2011). Quality control and preprocessing of metagenomic datasets. Bioinformatics, 27(6), 863–864. http://doi.org/10.1093/bioinformatics/btr026

Schnyder, E., Bodelier, P., Hartmann, M., Henneberger, R., & Niklaus, P. (2018). Positive diversity-functioning relationships in model communities of methanotrophic bacteria. Ecology, 99(3), 714–723. http://doi.org/10.1002/ecy.2138

Sierocinski, P., Bayer, F., Melia, G. Y., Großkopf, T., Alston, M., Swarbreck, D., … Buckling, A. (2018). Biodiversity – function relationships in methanogenic communities. Molecular Ecology, 27, 4641–4651. http://doi.org/10.1111/mec.14895

Sierocinski, P., Milferstedt, K., Bayer, F., Großkopf, T., Alston, M., Bastkowski, S., … Buckling, A. (2017). A Single Community Dominates Structure and Function of a Mixture of Multiple Methanogenic. Current Biology, 3390–3395.

Sjögersten, S., Black, C. R., Evers, S., Hoyos-Santillan, J., Wright, E. L., & Turner, B. (2014). Tropical wetlands: A missing link in the global carbon cycle? Global Biogeochemical Cycles, 28, 1371–1386. http://doi.org/10.1002/2014GB004844.Received

Steinberg, L. M., & Regan, J. M. (2008). Phylogenetic comparison of the methanogenic communities from an acidic, oligotrophic fen and an anaerobic digester treating municipal wastewater sludge. Applied and Environmental Microbiology, 74(21), 6663–71. http://doi.org/10.1128/AEM.00553-08

Tathy, J. P., Cros, B., Delmas, R. A., Marenco, A., Servant, J., & Labat, M. (1992). Methane emission from flooded forest in central Africa. Journal of Geophysical Research: Atmospheres, 97(D6), 6159–6168. http://doi.org/10.1029/90JD02555

Tilman, D., & Downing, J. A. (1994). Biodiversity and stability in grasslands. Nature, 367, 363–365.

Tilman, D., Wedin, D., & Knops, J. (1996). Productivity and sustainability influenced by biodiversity in grassland ecosystems. Nature, 379, 718. Retrieved from http://dx.doi.org/10.1038/379718a0

Wang, Q., Garrity, G. M., Tiedje, J. M., & Cole, J. R. (2007). Naive Bayesian Classifier for Rapid Assignment of rRNA Sequences into the New Bacterial Taxonomy. Applied and Environmental Microbiology, 73(16), 5261–5267. http://doi.org/10.1128/AEM.00062-07

White, F. (1979). The Guineo-Congolian Region and Its Relationships to Other Phytochoria. Bulletin Du Jardin Botanique National de Belgique/Bulletin van de National Plantentuin van Belgie, 49(1/2), 11–55. http://doi.org/10.2307/3667815

Wickham, H. (2009). ggplot2: elegant graphics for data analysis. Springer Science & Business Media.

Zhang, J., Kobert, K., Flouri, T., & Stamatakis, A. (2014). PEAR: a fast and accurate Illumina Paired-End reAd mergeR. Bioinformatics, 30(5), 614–620. http://doi.org/10.1093/bioinformatics/btt593

