## Supplemental Materials for "Community structure – ecosystem function relationships in the Congo Basin methane cycle depend on the physiological scale of function" (from another version of this preprint: version 1, 2019-05-17)

SUPPLEMENTARY MATERIALS

**Supplementary Table 1:** Methanogen and methanotroph genera included in the study. Note that multiple ASV-level taxa could come from within each genus.

| Functional group | Taxonomic Hierarchy |
|---|---|
| Methanogen | Archaea;Euryarchaeota;Methanobacteria;Methanobacteriales;Methanobacteriaceae;Methanobacterium; |
| Methanogen | Archaea;Euryarchaeota;Methanomicrobia;Methanocellales;Methanocellaceae;Methanocella; |
| Methanogen | Archaea;Euryarchaeota;Methanomicrobia;Methanomicrobiales;Methanoregulaceae;Methanolinea; |
| Methanogen | Archaea;Euryarchaeota;Methanomicrobia;Methanomicrobiales;Methanoregulaceae;Methanoregula; |
| Methanogen | Archaea;Euryarchaeota;Methanomicrobia;Methanomicrobiales;Methanoregulaceae;Methanosphaerula; |
| Methanogen | Archaea;Euryarchaeota;Methanomicrobia;Methanomicrobiales;Methanospirillaceae;Methanospirillum; |
| Methanogen | Archaea;Euryarchaeota;Methanomicrobia;Methanosarcinales;Methanosarcinaceae;Methanohalobium; |
| Methanogen | Archaea;Euryarchaeota;Methanomicrobia;Methanosarcinales;Methanosarcinaceae;Methanosalsum; |
| Methanogen | Archaea;Euryarchaeota;Methanomicrobia;Methanosarcinales;Methanosarcinaceae;Methanosarcina; |
| Methanogen | Archaea;Euryarchaeota;Methanomicrobia;Methanosarcinales;Methanotrichaceae;Methanothrix; |
| Methanogen | Archaea;Euryarchaeota;Methanomicrobia;Methanosarcinales;Methermicoccaceae;Methermicoccus; |
| Methanogen | Archaea;Euryarchaeota;Thermoplasmata;Methanomassiliicoccales;Methanomassiliicoccaceae;Methanomassiliicoccus |
| Methanotroph | Bacteria;Proteobacteria;Alphaproteobacteria;Rhizobiales;Beijerinckiaceae;Methylocapsa; |
| Methanotroph | Bacteria;Proteobacteria;Alphaproteobacteria;Rhizobiales;Beijerinckiaceae;Methylocella; |
| Methanotroph | Bacteria;Proteobacteria;Alphaproteobacteria;Rhizobiales;Beijerinckiaceae;Methyloferula; |
| Methanotroph | Bacteria;Proteobacteria;Alphaproteobacteria;Rhizobiales;Beijerinckiaceae;Methylorosula; |
| Methanotroph | Bacteria;Proteobacteria;Alphaproteobacteria;Rhizobiales;Beijerinckiaceae;Methylovirgula; |
| Methanotroph | Bacteria;Proteobacteria;Alphaproteobacteria;Rhizobiales;Methylocystaceae;Methylocystis; |
| Methanotroph | Bacteria;Proteobacteria;Alphaproteobacteria;Rhizobiales;Methylocystaceae;Methylosinus; |
| Methanotroph | Bacteria;Proteobacteria;Gammaproteobacteria;Methylococcales;Methylococcaceae;Methylobacter; |
| Methanotroph | Bacteria;Proteobacteria;Gammaproteobacteria;Methylococcales;Methylococcaceae;Methylocaldum; |
| Methanotroph | Bacteria;Proteobacteria;Gammaproteobacteria;Methylococcales;Methylococcaceae;Methylococcus; |
| Methanotroph | Bacteria;Proteobacteria;Gammaproteobacteria;Methylococcales;Methylococcaceae;Methylogaea; |

| | |
|---|---|
| Methanotroph | Bacteria;Proteobacteria;Gammaproteobacteria;Methylococcales;Methylococcaceae;Methylohalobius; |
| Methanotroph | Bacteria;Proteobacteria;Gammaproteobacteria;Methylococcales;Methylococcaceae;Methylomarinovum; |
| Methanotroph | Bacteria;Proteobacteria;Gammaproteobacteria;Methylococcales;Methylococcaceae;Methylomarinum; |
| Methanotroph | Bacteria;Proteobacteria;Gammaproteobacteria;Methylococcales;Methylococcaceae;Methylomicrobium; |
| Methanotroph | Bacteria;Proteobacteria;Gammaproteobacteria;Methylococcales;Methylococcaceae;Methylomonas; |
| Methanotroph | Bacteria;Proteobacteria;Gammaproteobacteria;Methylococcales;Methylococcaceae;Methyloparacoccus; |
| Methanotroph | Bacteria;Proteobacteria;Gammaproteobacteria;Methylococcales;Methylococcaceae;Methyloprofundus; |
| Methanotroph | Bacteria;Proteobacteria;Gammaproteobacteria;Methylococcales;Methylococcaceae;Methylosarcina; |
| Methanotroph | Bacteria;Proteobacteria;Gammaproteobacteria;Methylococcales;Methylococcaceae;Methylosoma; |
| Methanotroph | Bacteria;Proteobacteria;Gammaproteobacteria;Methylococcales;Methylococcaceae;Methylosphaera; |
| Methanotroph | Bacteria;Proteobacteria;Gammaproteobacteria;Methylococcales;Methylococcaceae;Methylovulum; |

**Supplementary Table 2:** Characterization of the physical and chemical variables measured across all sites. Averages with standard errors shown. ET = ecosystem type; WT = water table; T = temperature; $MC_a$ = moisture content above the water table.

| ET | pH | WT (± cm) | Soil T (°C) | % Total N | % Organic C | % Clay | % Sand | % Silt | $MC_a$ (%) |
|---|---|---|---|---|---|---|---|---|---|
| Open Peatland | 3.8 ± 0.02 (n=4) | -11.3 ± 3.1 (n=4) | 29.6 ± 0.3 (n=4) | 1.6 ± 0.01 (n=3) | 39.6 ± 0.9 (n=3) | 0 | 0 | 0 | 89.2 ± 0.9 (n=3) |
| Forested Peatland | 3.6 ± 0.1 (n=3) | -22.9 ± 3.9 (n=3) | 26.6 ± 0.1 (n=3) | 1.2 ± 0.1 (n=2) | 45.7 ± 1.6 (n=2) | 0 | 0 | 0 | 74.7 ± 3.1 (n=3) |
| Forested | 6.1 ± | -6.9 ± | 26.9 ± | 1.3 ± 0.7 | 24.7 ± | 1.2 | 95.1 | 3.7 | 60.1 ± |

| | | | | | | | | | |
|---|---|---|---|---|---|---|---|---|---|
| MSW | 0.1 (n=5) | 1.5 (n=5) | 0.2 (n=5) | (n=2) | 16.0 (n=2) | | | | 17.9 (n=3) |
| Forested MSW | 5.0 ± 0.1 (n=4) | -1.7 ± 7.2 (n=4) | 26.1 ± 0.1 (n=4) | 0.4 ± 0.2 (n=2) | 5.5 ± 2.8 (n=2) | 3.2 | 84.2 | 12.6 | 49.7 ± 3.9 (n=3) |
| Forested MSW | 4.7 ± 0.3 (n=5) | -5.1 ± 3.4 (n=5) | 26.9 ± 0.2 (n=5) | 0.5 ± 0.4 (n=2) | 9.9 ± 9.3 (n=2) | 2.3 | 80.1 | 17.6 | 49.1 ± 13.3 (n=3) |
| Forested MSW | 4.8 ± 0.1 (n=6) | -10.9 ± 5.3 (n=6) | 27.0 ± 0.1 (n=6) | 0.2 ± 0.04 (n=3) | 5.1 ± 2.5 (n=3) | 1.2 | 75.3 | 23.5 | 31.9 ± 10.4 (n=3) |
| Seasonal MSW | 4.2 ± 0.1 (n=3) | N/A | 26.4 ± 0.1 (n=6) | 0.3 ± 0.05 (n=3) | 3.6 ± 0.7 (n=3) | 1.5 | 79.6 | 18.9 | 22.9 ± 2.1 (n=3) |
| Seasonal MSW | 4.1 ± 0.1 (n=3) | N/A | 26.7 ± 0.1 (n=6) | 1.2 ± 0.2 (n=3) | 23.6 ± 4.2 (n=3) | 1.7 | 98.3 | 0 | 53.5 ± 3.7 (n=3) |
| Upland Grassland | 4.3 ± 0.03 (n=3) | N/A | 29.6 ± 0.11 (n=6) | 0.1 ± 0.07 (n=3) | 3.7 ± 2.9 (n=3) | 0.4 | 99.2 | 0.4 | 1.7 ± 0.4 (n=3) |
| Upland Grassland (with termite mounds) | 4.7 ± 0.2 (n=3) | N/A | 28.0 ± 0.03 (n=6) | 0.04 ± 0.01 (n=3) | 0.8 ± 0.1 (n=3) | 1.1 | 98.1 | 0.8 | 4.6 ± 0.5 (n=3) |
| Upland Forest | 3.9 ± 0.03 (n=3) | N/A | 26.5 ± 0.3 (n=5) | 0.2 ± 0.02 (n=3) | 2.1 ± 0.2 (n=3) | 3.6 | 84.2 | 12.2 | 11.1 ± 1.2 (n=3) |
| Upland Forest | 4.1 ± 0.1 (n=3) | N/A | 26.7 ± 0.1 (n=6) | 0.2 ± 0.01 (n=3) | 2.0 ± 0.3 (n=3) | 4.1 | 66.9 | 29.0 | 12.9 ± 0.1 (n=3) |
| Upland Forest | 4.0 ± 0.06 (n=3) | N/A | 27.0 ± 0.1 (n=6) | 0.4 ± 0.2 (n=3) | 7.3 ± 4.0 (n=3) | 0 | 89.3 | 10.7 | 23.5 ± 7.3 (n=3) |
| Upland Plantatio | 4.0 ± 0.1 | N/A | 31.0 ± 0.3 | 0.1 ± 0.04 (n=3) | 1.3 ± 0.5 (n=3) | 8.1 | 66.0 | 25.9 | 12.1 ± 1.2 |

| | | | | | | | | | |
|---|---|---|---|---|---|---|---|---|---|
| n | (n=3) | | (n=5) | | | | | | (n=3) |
| Upland Ab. Plantatio n | 4.1 ± 0.03 (n=3) | N/A | 27.0 ± 0.3 (n=6) | 0.1 ± 0.03 (n=3) | 1.8 ± 0.6 (n=3) | 3.3 | 95.9 | 0.8 | 6.2 ± 1.6 (n=3) |

**Supplementary Table 3**: The compositional correlates of high-affinity methanotrophy. Inference method refers to DNA- or RNA-based 16S rRNA gene profiling. Taxonomic assignments to the finest levels are presented.

| Inference Method | Taxonomy |
|---|---|
| DNA | |
| | Archaea\| Thaumarchaeota\| Nitrososphaerales\| Nitrososphaeraceae\| Nitrososphaera |
| | Bacteria\| Acidobacteria\| Acidobacteria_Gp6\| Gp6 |
| | Bacteria\| Proteobacteria\| Alphaproteobacteria\| Rhizobiales\| Roseiarcaceae\| Roseiarcus |
| | Bacteria\| Acidobacteria\| Acidobacteria_Gp6\| Gp6 |
| | Bacteria\| Acidobacteria\| Acidobacteria_Gp2\| Gp2 |
| | Bacteria\| Acidobacteria\| Acidobacteria_Gp2\| Gp2 |
| | Bacteria\| Actinobacteria\| Actinobacteria\| Actinomycetales\| Thermomonosporaceae |
| | Bacteria\| Acidobacteria\| Acidobacteria_Gp2\| Gp2 |
| | Bacteria\| Acidobacteria\| Acidobacteria_Gp2\| Gp2 |
| | Bacteria\| Synergistetes\| Synergistia\| Synergistales\| Synergistaceae\| Aminiphilus |
| | Bacteria\| Proteobacteria\| Alphaproteobacteria\| Rhodospirillales\| Acetobacteraceae\| Acidisoma |
| RNA | |
| | Bacteria\| Proteobacteria\| Alphaproteobacteria\| Rhizobiales\| Beijerinckiaceae\| Beijerinckia |
| | Bacteria\| Actinobacteria\| Actinobacteria\| Actinomycetales\| Thermomonosporaceae |
| | Bacteria\| Proteobacteria\| Deltaproteobacteria\| Deltaproteobacteria_incertae_sedis\| Deferrisoma |
| | Bacteria\| Proteobacteria\| Alphaproteobacteria\| Rhodospirillales\| Rhodospirillaceae\| Nitrospirillum |
| | Bacteria\| Acidobacteria\| Acidobacteria_Gp1\| Gp1 |
